# Neuronal secretome from bipolar patient-derived neurons alters network function and contains candidate biomarkers for diagnosis and lithium response

**DOI:** 10.64898/2026.05.11.724377

**Authors:** Alessia Pietrantonio, Charles-Étienne Castonguay, Daniel Rochefort, Camila Tiefensee-Ribeiro, Yumin Liu, Malak Abuzgaya, Isabella Pietrantonio, Ziqi Yu, Martin Alda, Guy A. Rouleau, Austen J. Milnerwood, Anouar Khayachi

**Author notes:** Equal Contribution (Guy A. Rouleau, Austen J. Milnerwood, Anouar Khayachi). Corresponding authors (G.A. Rouleau), (A.J. Milnerwood), (A. Khayachi).

## Abstract

Delayed diagnosis and treatment are a major burden to patients with bipolar disorder. While lithium is the most effective treatment against mania, depressive episodes, and suicide, only 30% of patients respond to it fully. Currently there are no reliable methods to predict lithium responsiveness. To address these challenges, we aimed to identify potential diagnostic and treatment response biomarkers for BD, in addition furthering understanding of BD pathophysiology. Here, we leveraged human induced pluripotent stem cell (hiPSC) derived neurons from lithium responsive (LR), lithium non-responsive (LNR), and healthy age-matched controls (CTL). We found extracellular vesicle (EV) cargos from hiPSC-derived neurons are indicative of disease state and treatment-response. Unbiased proteomic and miRNA profiling identified 10 proteins and 13 miRNAs that were differentially expressed in BD EVs relative to CTL, as well as distinct molecular signatures separating LR ad LNR groups. These differences converged on pathways related to synaptic function, neurotrophic signalling, and cellular stress responses. Additionally, we found the BD neuronal secretome alters activity in non-BD neuronal networks. Chronic treatment of CTL cultures with BD neuron-conditioned media modified the proportion of active neurons and the frequency and amplitude of calcium transients in individual neurons. We demonstrate that neuronal EVs contain molecular signatures of disease state and treatment response in BD and identify the BD secretome as an active regulator of neuronal network homeostasis. This study provides novel insights into the pathophysiology of BD and candidate biomarkers for personalized BD diagnosis and treatment selection.

## Introduction

Bipolar disorder (BD) is a severe mood disorder with a global prevalence of 1-1.5%^1^, characterized by recurrent episodes of mania and depression^2^. It is highly heritable (>75%)^3^, but genetically complex; genome wide association studies have identified 298 risk loci but no single causal variant^4^. Overall, identified GWAS loci implicate genes involved in synaptic biology^5^, and variants in genes encoding calcium, sodium, and potassium channels suggest ion channel dysfunction is a BD disease mechanism^5^. Despite many advances, the biological basis of BD remains incompletely understood, in part due to clinical and genetic heterogeneity.

Diagnosis of BD is challenging due to substantial symptom overlap with other mood disorders^6^. Thus, misdiagnosis is common and long delays to correct diagnosis lead to a more severe prognosis and a high suicide rate^7^. Treatment is often selected by trial-and error, further delaying effective mood stabilization. These facts highlight a need for biomarkers to guide diagnosis and treatment stratification. Lithium (Li) is the most effective mood stabilizer for long-term relapse prevention, yet only ∼30% of BD patients respond robustly, separating BD patients into two groups: lithium responders (LR) and lithium non-responders (LNR).

With the development of induced pluripotent stem cell (iPSC) technology, it is possible to study the effect of Li on BD patient-derived neurons. Recently, we and others demonstrated a hyperexcitability phenotype in BD patient-derived neurons that is reversible by lithium in neurons from LR, but not LNR, patients^8–10^. However, it remains unclear what causes the hyperexcitability of BD iPSC-derived neurons, and how this translates to BD pathophysiology.

Nevertheless, these advances can be leveraged to try to improve BD diagnosis and treatment, and thus the quality of life of patients, by identifying relevant biomarkers. Approaches to diagnostic biomarker discovery in other scenarios has seen the emergence of extracellular vesicles (EV) as a promising accessible source^11^. EVs are released by most cell types and carry proteins, lipids, and nucleic acids reflective of their cell of origin^12, 13^. Importantly, brain-derived EVs cross the blood–brain barrier and are detected in peripheral biofluids, enabling access to neuronal molecular signatures^12^. Beyond diagnostic potential, EVs are increasingly recognized as mediators of intercellular communication, capable of influencing recipient cell function^14–17^.

While prior studies have identified EV-associated miRNAs in BD patient biofluids^18^, relatively few have examined brain-derived EVs^19^, which may more directly reflect disease-relevant processes. Notably, differences in clinical relevance between brain-derived and peripheral EV signatures highlight the importance of cellular context in biomarker discovery^20^.

Here, we isolated EVs from hiPSC-derived forebrain neurons from CTL, LR, and LNR to identify candidate biomarkers and investigate disease mechanisms. Using proteomic and miRNA profiling, we identified distinct EV signatures associated with BD and lithium response pointing to molecular pathways related to synaptic function, neurotrophic signaling, and cellular stress responses. We also demonstrated that the BD secretome modulates neuronal network activity, indicating that EV-associated factors are not only biomarkers but also active contributors to network dysfunction. Together, these findings position neuronal EVs as both informative readouts and active mediators of disease-relevant processes, providing a framework to link patient-specific cellular dysfunction to network-level phenotypes and to advance biomarker discovery and mechanistic stratification in bipolar disorder.

## Methods

### HeLa and U2OS cell culture

HeLa and U2OS cells were maintained in Dulbecco’s Modified Eagle Medium supplemented with 10% fetal bovine serum and 1% Penicillin/Streptomycin at 37°C with 55 CO2. Cells were passaged using trypsin-EDTA and used for EV collection at 80% confluency following at least 2 passages post-thaw.

### hiPSC generation and culture

Patient-derived iPSC lines (Supp. Table S1) were generated from peripheral blood mononuclear cells or lymphocytes of LR, LNR, and age-matched controls. Lithium response was defined using the Alda scale for Li responsiveness (LR: 9-10/10, LNR: 0-3/10)^21–23^. Reprogramming was performed as previously described^8, 9^. hiPSCs were maintained in mTeSR^TM^1 (STEMCELL technologies) on Matrigel (Corning) and passaged at least twice prior to neural induction.

### Neural induction and differentiation

#### Embryoid body (EB) protocol

On day 0, a single cell suspension of IPSCs was generated using Gentle Cell dissociation reagent (STEMCELL technologies). In an ultra-low attachment 96-well plate, 10,000 cells/well were plated in STEMdiff^TM^ neural induction medium supplemented with SMADi and 10uM of Y-27632 (Selleckchem). ¾ medium changes were performed on until day 5 where EBs were replated on poly-L-ornithine (PLO, Sigma) and laminin (Fisher) coated 6-well plates. Full media changes were performed from days 6-11, and neural induction efficiency was determined to be greater than 75%. On day 12, neural rosette selection was performed using the STEMdiff^TM^ Neural Rosette Selection Reagent and replated into PLO/laminin coated 6-well plates. Full medium changes were performed from days 13-19. NPCs were passaged with Gentle Cell Dissociation Reagent at 80-90% confluence, typically between days 17-19. NPCs were cultured in STEMdiff^TM^ Neural Progenitor Medium and were passaged twice before proceeding to neural differentiation.

#### Monolayer protocol

On day 0, a single cell suspension of iPSCs was generated using Gentle Cell dissociation reagent. 2 x 10^6^ cells/well were plated in STEMdiff^TM^ neural induction medium supplemented with SMADi and 10uM of Y-27632 in a PLO/laminin coated 6-well plate. Daily medium changes were performed until cultures were ready to be passaged (typically between days 6-9). Cells were lifted using ACCUTASE^TM^ (STEMCELL technologies) and passaged into PLO/laminin coated 6-well plates. After approximately 7 days of culture, this process was repeated before proceeding to the first passage using STEMdiff^TM^ Neural Progenitor Medium as described in the EB protocol above.

### Forebrain neuron differentiation

NPCs were lifted using ACCUTASE^TM^, and differentiated in PLO/laminin coated 10cm dishes (for EV preparation) or 12mm glass coverslips in 24 well plates (for calcium imaging experiments) in final differentiation media (BrainPhys^TM^ (STEMCELL Technologies), Glutamax (Fisher), N2 (Fisher), NeuroCult^TM^SM1 Neuronal Supplement (STEMCELL technologies), 200nM ascorbic acid (STEMCELL Technologies), 500ug/mL cyclicAMP (SIGMA), 20ng/mL brain-derived neurotrophic factor (GIBCO), 10ng/mL Wnt3a (R&D Systems), and 1ug/mL laminin (GIBCO 23017015)). Final differentiation media changes were performed every other day for 2 weeks. After 2 weeks, STEMdiff^TM^ Forebrain Neuron Maturation kit (STEMCELL technologies) was used to change media twice per week for 4-6 weeks.

### EV isolation

We used a 8% polyethylene glycol (PEG) precipitation + UC wash method based on a protocol by Rider et al.^24^. Conditioned neuronal media was collected 3x from 10cm dishes 28-32 days post-differentiation. Media was centrifuged at 500g for 5 minutes, the supernatant filtered through a 220nm filter, and subsequently centrifuged at 2000 xg for 30 minutes. The supernatants were then mixed with an equal volume of 2X PEG stock solution (16% m/v PEG 6000 (PB0432, BioBasic) in 1M NaCl in filtered water) and rotated overnight at 4 °C. Samples were centrifuged at 4000rpm at 4°C for 1 hour (Thermo Fisher, MegaFuge^TM^ 40R), and the pellet resuspended in 500uL filtered PBS. The suspension was ultracentrifuged (Beckman Coulter, Optima^TM^ MAX) in polycarbonate centrifuge tubes (Beckman Coulter, 343776) at 108,000 xg in a fixed angle rotor (Beckman Coulter, TLA-120.1) for 70 minutes at 4 °C. The resulting EV pellet was then resuspended in 50uL filtered PBS (for proteomics and NTA), 50uL TRizol (for miRNA sequencing), or 50uL of lysis buffer (for western blot) and stored at -80 °C until use. For EV isolation from U2OS and HeLa cells, media was collected once from a single 10cm dish at 80% confluency. For NPCs, media was collected once from two T75 flasks.

### Electron microscopy

Data were collected on the FEI Tecnai G2 Spirit Twin 120kV Cryo-Transmission electron microscope (TEM) located at the Facility for Electron Microscopy Research (FEMR) at McGill University. EVs were fixed with 2.5% glutaraldehyde in 0.1M sodium cacodylate buffer.

#### Sample preparation

The negative staining protocol and grid preparation was performed following FEMR guidelines. Carbon-coated 200-mesh copper TEM grids were glow-discharged using the Pelco easiGlow^TM^ Discharge Cleaning System. 5uL of EV sample incubated on the TEM grid for 10 minutes. The grid was then transferred onto drops of 0.2M glycine solution, then washed dH2O. Next, the grid was stained using filtered 2% uranyl-acetate and air dried at room temperature for a minimum of 1 hour prior to imaging by TEM (AMT NanoSprint15 MK2 CMOS camera) at 120kV.

### Nanoparticle tracking analysis (NTA)

Collected media was processed via the 8%PEG+UC protocol and frozen at -80C. An equal volume of STEMdiff^TM^ Forebrain Neuron Maturation kit (STEMCELL technologies) media was collected as a negative control. Samples were brought to the Center for Applied Nanomedicine at the McGill University Health Center to be measured via NTA using The ZetaView® PMX-120 (Particle Metrix) with a highly sensitive 640 by 680 pixels CMOS camera. 11 positions were recorded throughout each EV sample.

### Western blot

Lysis buffer (10mM Tris HCl, 10mM EDTA, 150mM NaCl, 1% triton X100, 0.1% SDS, in distilled water) was added to final EV pellets and cell lysates. The samples were then rotated at 4°C for 30 minutes, followed by probe sonication. Protein quantification was done using the Pierce BCA^TM^ assay (ThermoFisher 23255) for cell lysate, and the Pierce Micro BCA^TM^ assay (ThermoFisher 23235) for EV samples. Samples were then denatured using 4x NuPage LDS sample buffer (Invitrogen NP0008) and β-mercaptoethanol (BME) for a final concentration of 1X LDS and 5% BME. After heating for 5 minutes at 95 °C, 10ug (for EV validation) and 25ug (for NTA characterization) of each sample was loaded into hand-cast 10% Bis-tris gels and run at 70V, followed by 100V in a Mini-PROTEAN^®^ Tetra cell (BioRad) gel electrophoresis tank.

Proteins were transferred to methanol-activated Immobilon-FL PVDF membranes (Millipore IPFL00010) for 900 minutes at 25V and at 4 °C. Total protein was observed using the Revert^TM^ 700 Total Protein Stain Kit (LICORbio) prior to blocking for 1 hour in 5% BSA in TBS+1% tween 20 (TBS-T). Primary antibodies incubated overnight in 3% BSA in TBS-T at 4 °C, then washed 3X for 15 minutes with TBS-T. Membranes were incubated with appropriate fluorescent LI-COR secondary antibodies in 3% BSA in TBS-T for 1 hour at room temperature, before repeating TBS-T washes, and imaging on the LI-COR Odyssey Infrared imaging system (LI-COR). The imaged membranes were background subtracted, and the resulting bands were analyzed using Fiji software. Primary antibodies: ALIX (1/1000, Abcam, ab117600), TSG 101 (1/1000, Abcam, ab30871), L1CAM (1/5000, Abcam, ab24345), HSP90α (F-2) (1/500, Santa Cruz Biotechnology, sc-515081), GFAP (1/5000, Biotech, NB300-141), TUJ1 (1/1000, Sigma, T8660), PAX6 (1/2000, Biolegend, 901301), GAPDH (1/2000, Invitrogen, MA5-15738).

### Proteomics

Conditioned media from 3×10cm dishes per patient-derived neuronal line was collected at div28-32 after differentiation. An equal volume of STEMdiff^TM^ Forebrain Neuron Maturation kit (STEMCELL technologies) media was also collected as a negative control. Collected media was processed via the 8%PEG+UC protocol and frozen at -80C. Samples were brought to the proteomic platform at the Institute for Research in Immunology and Cancer located at Université de Montréal. Samples were trypsinized and underwent unbiased label-free quantitative proteomics using data independent acquisition (DIA) liquid chromatography tandem mass spectrometry (LC-MS/MS). All proteomic comparisons and statistics were performed using the scaffold DIA V4.2.1 software.

### RNA extraction

Samples were spiked-in with QIAseq miRNA Library QC Spike-ins (QIAGEN, 331535) during RNA extraction at a reduced concentration (1/1000) relative to manufacturer’s instructions to account for low total RNA input. Samples were then homogenized with QIAshredder (Qiagen, 79654) mini spin columns prior to RNA extraction. RNA extraction was done following manufacturer’s instructions for the Direct-zol RNA Microprep kit (ZYMO Research, R2060).

### cDNA library generation

cDNA libraries were generated following manufacturer’s instructions for the NEBNext^®^ Low-bias Small RNA Library prep kit (NEB, #E3420S) in combination with NEBNext^®^ LV Unique Dual Index Primers Set 2A and 2B (NEB, #E3390S, #E3392S). RNA input was based on quantifications using the Qubit^TM^ RNA HS Assay kit (Invitrogen, Q32852). Primers and adapters were diluted 1:2 based on quality control (QC) assessments. Bead cleanup of PCR amplification reaction was performed without size selection to maximize cDNA output.

### microRNA sequencing

Libraries were sequenced by Génome Québec on an Illumina HiSeq2000 platform using single end 100bp reads (SE100), with up to 400 million reads per run. Sequencing depth per sample was adjusted based on library QC assessments. Libraries with suboptimal QC metrics (low concentration or presence of dimers) were sequenced at a reduced pooling ratio (1:10) relative to higher-quality libraries which were sequenced at an equimolar ratio (1:1).

### microRNA sequencing analysis

MicroRNA sequencing reads were processed and quantified using miRge3 with MirGeneDB using the default settings. Differential expression analysis was performed using DESeq2 with a multifactor design to account for group, treatment, and their interaction. MiRNAs with counts ≥1 in at least six samples were retained, counts were normalized for library size, and statistical significance was determined using Wald tests with a false discovery rate threshold of 0.1. miRNA gene targets of differentially expressed miRNA’s were determined using the miRTarBase database (https://awi.cuhk.edu.cn/~miRTarBase/miRTarBase_2025/php/index.php) ^25^. Upregulated and downregulated miRNAs were assessed separately, and target genes were restricted to those experimentally validated by WB, qPCR, and reporter assay. Enrichr (https://maayanlab.cloud/Enrichr/) ^26^, a gene set search engine, was used to perform GO enrichment analysis on the gene targets identified using miRTarBase.

### Calcium Imaging

#### Viral Transduction

Coverslips (12mm) of approximately 85,000 iPSC-derived neurons were transduced by adeno-associated virus vectors, encoding GCaMP6f under a human synapsin promoter (Serotype: AAV2/PHP.eB; Canadian Neurophotonics Platform Viral Vector Core Facility (RRID:SCR_016477)) 9 days prior to recording.

#### Neuronal secretome treatment (*Fig. 6a*)

Conditioned media was collected from BD neurons (LR or LNR) that were differentiated in parallel to CTL neurons (same age post-differentiation). CTL neurons were treated with conditioned media containing the BD secretome from LR or LNR starting at 16 days post-differentiation, until recording. CTL neurons that did not receive treatment, received a half-media change as a vehicle control.

#### Optical Imaging

Coverslips were placed in a recording chamber on an Olympus BX51 upright microscope stage, perfused with artificial cerebrospinal fluid (aCSF; in mM: 125 NaCl, 3 KCl, 25 NaHCO_3_, 1.25 NaH_2_PO_4_, 2 MgCl_2_, 2 CaCl_2_, 10 glucose, pH 7.2-7.4, 300-310 mOsm) at 25-26 °C. Live-cell imaging of coverslips was performed at 34-42 days (5-6 weeks) post-neural differentiation with a 20x immersive objective. Bright field and GCaMP6f fluorescence (520nm) were stimulated by a LED (white light, and CoolLED) and visualized through a GFP bandpass filter and an EM-CCD camera (Andor iXon Ultra 897) controlled by Andor Solis software. Video trial acquisition was 800 frames of 512×512 pixel image fields, recorded at ∼3.7 Hz (∼3.6 minutes). Recordings of 3 independent image fields for each coverslip were acquired without treatment, followed by recording of the same 3 fields after switching to aCSF containing 2.5 mM of the weak K+ channel blocker 4-Aminopyridine (4-AP) to increase activity. Coverslips were perfused with aCSF+4-AP for 7 minutes prior to treated recordings. Brightfield images were taken of each field for reference during analysis and ROI selection. Cell soma were selected as regions of interest using Suite2P^27^, followed by quantification of calcium event frequency, amplitude, and area under the curve using a custom python script.

## Statistics

All statistical analyses were performed using GraphPad Prism 11 software (GraphPad software, Inc.). All data are represented as the mean +/- standard error of the mean (SEM). A p-value < 0.05 was considered significant. Specific statistical tests used are described in detail in the figure legend.

## Results

### Characterization of EV’s from iPSC-derived neurons

EVs were isolated using a combined polyethylene glycol precipitation and ultracentrifugation (PEG+UC) approach, selected for its balance of yield and purity ^24, 28, 29^. EV enrichment was confirmed by immunoblot detection of canonical markers (L1CAM, Alix, and TSG101) and absence of the non-vesicular protein HSP90α (Fig. 1a,b)^28^. Proteomic profiling further supported EV enrichment, with 97/100 and 100/100 of the top EV-associated proteins identified from ExoCarta (http://www.exocarta.org)^30^ and Vesiclepedia (http://www.microvesicles.org)^31^ databases, respectively (Fig. 1c).

**Figure 1:**
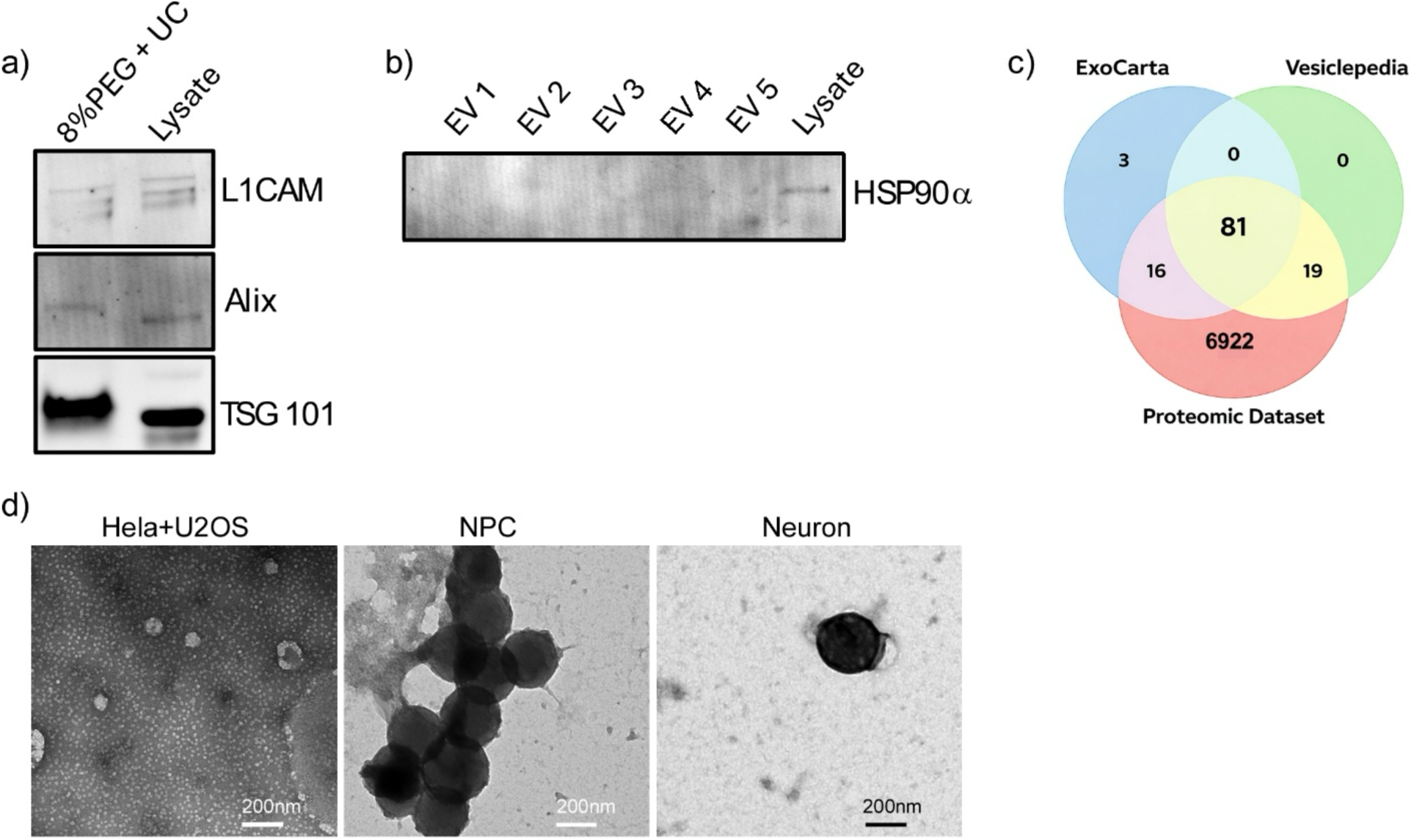
EV isolation and characterization. **a)** Western blot comparing the presence of canonical EV markers L1CAM (200 kDa), Alix (96 kDa), and TSG101 (49 kDa) between the 8% PEG + Ultracentrifugation (8% PEG + UC) technique and total cell lysate. **b)** Western blot targeting HSP90α (90kDa), a non-vesicular extracellular particle, from an n=5 EV samples compared to total cell lysate. **c)** Venn diagram comparing the top 100 EV proteins from two online EV databases (ExoCarta and Vesiclepedia [44,45]) to the proteomic dataset obtained from our isolated EVs (n=13). **d)** Representative transmission electron microscopy images of EVs extracted from pooled HeLa and U2OS cells (left), patient-derived NPCs (middle), and patient-derived neurons (right) stained with 2% uranyl-acetate.

The next step of characterization was to confirm the morphology, size distribution, and structural integrity of the isolated EVs. EVs were isolated from U2OS&HeLa cells, NPCs, and neurons and visualized by transmission electron microscopy (TEM). EVs from neurons and NPCs were circular, with a size of 50-200nm in diameter (Fig. 1d), consistent with the expected size range of our preparation, which includes a 220nm syringe filtration step. EVs isolated from highly secretory U2OS and HeLa cells were smaller and more abundant than those secreted from neurons (Fig. 1d). Neuron-derived EVs exhibited greater electron density compared to NPCs and non-neuronal cell lines, suggesting differences in vesicular cargo composition (Fig. 1d).

### Assessing differences in EV release between patient groups of iPSC-derived neurons

We next assessed EV release from CTL (n=3), LR (n=5), and LNR (n=4) neurons using nanoparticle tracking analysis (NTA). We first ensured there were no variations in gross cellular composition across cultures. Immunoblot analysis using neuronal (TUJ1), astrocytic (GFAP), and NPC (PAX6) markers demonstrated comparable cellular composition between CTL and BD-derived cultures normalized to GAPDH (Fig. 2a-b) and to total protein (Supp. Fig. S2a-b). NTA identified a significant interaction between vesicle size and bipolar status (Fig. 2c). LR neurons exhibited increased EV release relative to CTL across a defined size range (77.5-212.5nm) and in total particle concentration (Fig. 2 c&d). Results were consistent between GAPDH-normalized (Fig. 2c-d), raw (Supp. Fig. S2c-d), and total protein-normalized values (Supp. Fig. S2e-f).

**Figure 2:**
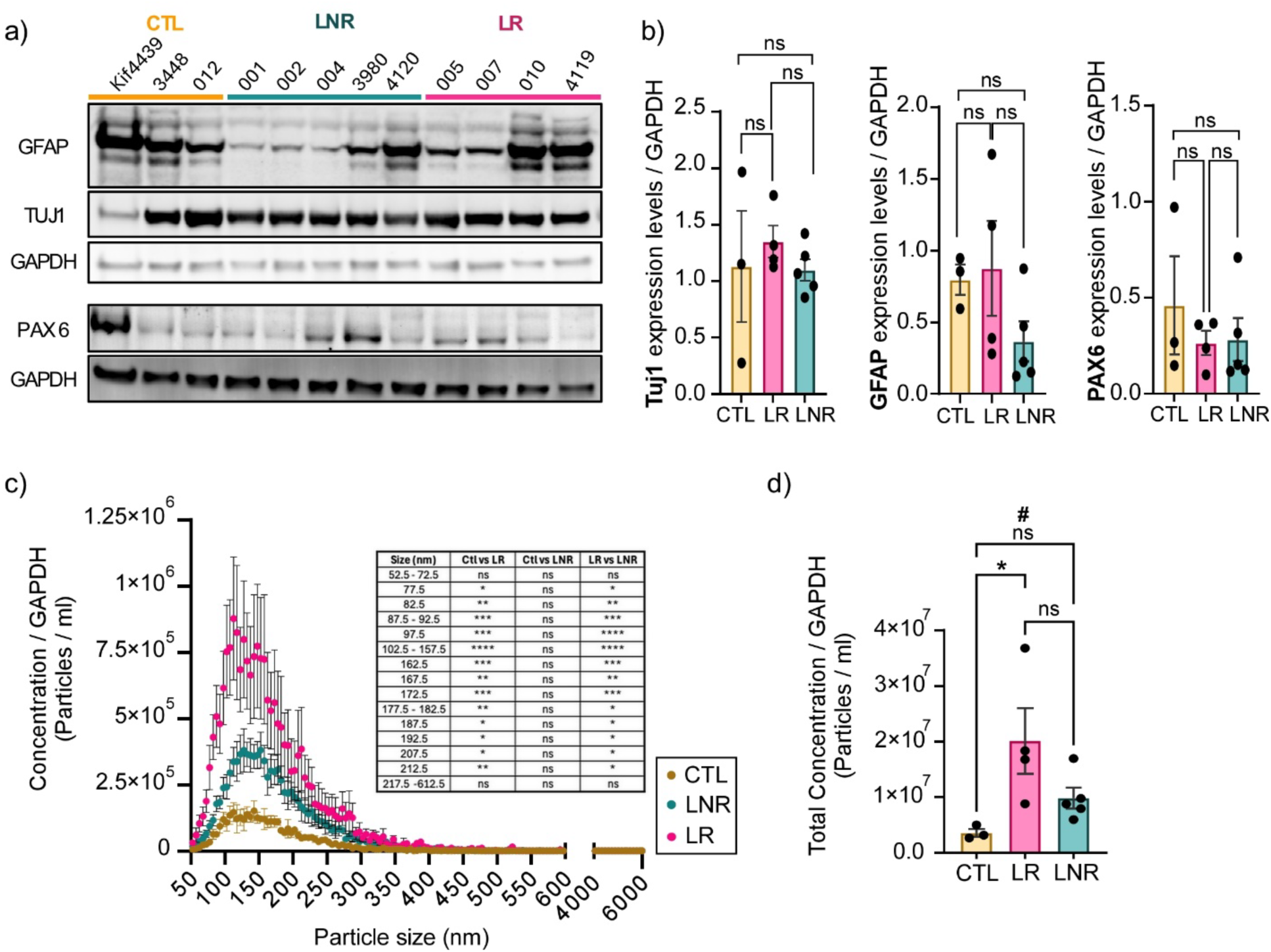
Characterization of EV outputs from patient-derived neurons. **a)** Western blot comparing cell-type composition between CTL (n=3), LNR (n=5), and LR (n=4) iPSC-derived neurons used for nanoparticle tracking analysis (NTA) experiments. Astrocyte, neuron, and NPC composition were assessed using GFAP (50 kDa), TUJ1 (50 kDa), and PAX6 (47 kDa) respectively, with GAPDH (37 kDa) as a loading control. **b)** Quantification of **(a)** normalized to GAPDH. **c)** NTA of isolated EVs from CTL (yellow, n=3), LR (pink, n=4), and LNR (turquoise, n=5) iPSC-derived neurons normalized to GAPDH. **d)** Total concentration of EVs isolated from CTL, LR, and LNR iPSC-derived neurons normalized to GAPDH. *Data shown are mean ±SEM. Statistics: One-way ANOVA with Tukey’s multiple comparisons test **(b)**, two-way ANOVA with Holm-Šídák’s multiple comparisons test **(c)** and one-way ANOVA with Šídák’s multiple comparisons test **(d)**. **\***p<0.0332, **p<0.0021, ***p<0.0002, ****p<0.0001. # Welch’s t-test of CTL vs LNR was statistically significant (p=0.0259)*.

Although LNR release >2 fold more EVs than CTL (Fig. 2c&d), the difference did not reach significance by *post hoc* testing correcting for multiple comparisons (Fig. 2d, *p=0.2764*); however, a direct comparison between CTL and LNR shows LNR neurons also secrete significantly more EVs than CTRL (Welch’s t-test *p=0.0259*, Fig. 2d). In addition, the concentration of EVs released from LR neurons across a size range of 77.5nm - 212.5nm was significantly increased relative to LNR (Fig. 2c), but there was no difference between LR & LNR when comparing total EV concentration (Fig. 2d). The data indicate that BD neurons exhibit increased EV secretion, particularly within the “small” EV population.

### EV proteomics reveals BD and lithium response associated signatures

Identifying proteins that are differentially expressed in EVs isolated from CTL, LR, and LNR neurons can not only allow us to better understand the intrinsic neuronal differences between these patient groups, but also detect protein biomarkers associated with BD. We performed unbiased label-free quantitative proteomics using data independent acquisition (DIA) liquid chromatography tandem mass spectrometry (LC-MS/MS). EVs were isolated from CTL (n=4), LR (n=4), and LNR (n=5) patient-derived neurons. Comparisons were performed between CTL and BD EVs to identify candidate BD-associated biomarkers, and between LR and LNR to investigate the biological differences associated with Li responsiveness.

From a total of 7047 identified proteins, statistical analyses revealed 10 and 28 differentially expressed in CTL vs BD and LR vs LNR EVs, respectively (Fig. 3a-d). Differentially expressed proteins in the CTL vs BD comparison that reached statistical significance (p<0.05), were 2-5-fold enriched in BD EVs relative to CTL (Fig. 3a-b). These were grouped into functional categories based on their roles in synaptic plasticity (NRXN1, SYT11, KIDINS220), synapse formation (THBS1, APLP2, ISLR2, DNER), neurotrophic signalling and survival (CPE, KIDINS220), and lipid metabolism (LIPG, LPL) (Fig. 3b).

**Figure 3:**
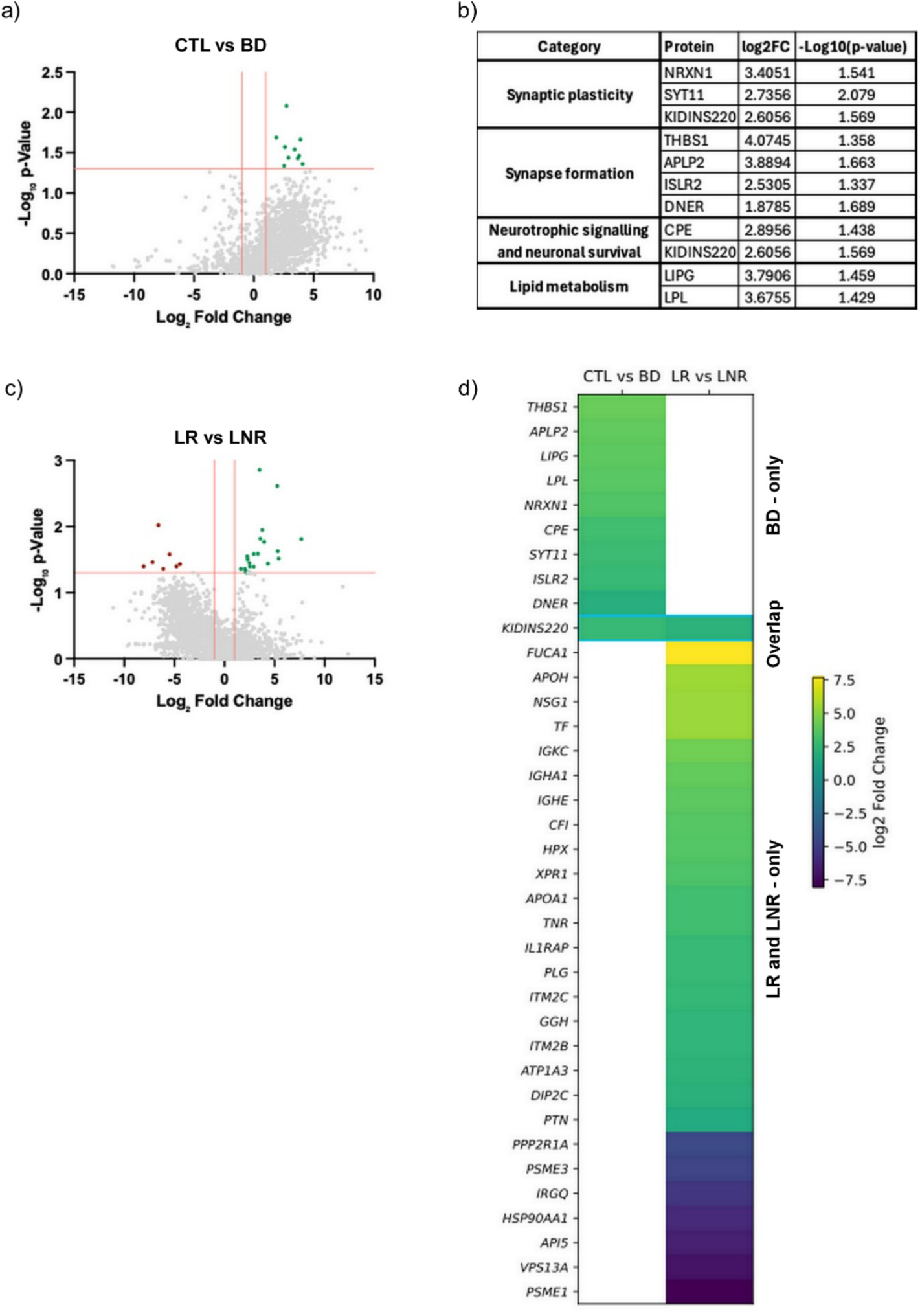
Proteomic profiling of patient-derived EVs. **a)** Volcano plot comparing differential protein expression in EVs from CTL (n=4) vs BD (n=9) neuronal cultures. *X-intercept: Log_2_FC= ±1. Y-intercept: –log_10_(P-value) = 1.3 (p = 0.05). Data points represent individual proteins. Points in grey are not significant, points in green are proteins enriched in BD EVs. **b***) Table summarizing differentially expressed proteins from **a)**, their functional category based on scientific literature, log_2_ fold change, and -log_10_(p-value). **c)** Same as in **a)**, but for EVs isolated from LR (n=4) vs LNR (n=5). *Points in green are proteins enriched in LR. Points in red are enriched in LNR.* **d)** Heatmap of differentially expressed proteins from **(a)** & **(c)**. 9 differentially expressed proteins are present in pooled BD EVs, 27 are differentially expressed between LR and LNR, and 1 is present in both conditions. *Statistics: Comparisons were t-test with p <0.05, and FDR < 1%*.

Notably, EVs from LR neurons had a >2 fold increase in ATP1A3 (Na^+^/K^+^-ATPase α3) (Fig. 3d), a neuron-enriched Na^+^/K^+^-ATPase subunit essential for maintaining membrane potential and regulating neuronal excitability^32^. Two additional proteins of interest enriched in LR EVs were KIDINS220 and pleiotrophin (PTN), both of which are involved in neurotrophic signalling, a pathway implicated elsewhere in Li’s therapeutic response^33^(Fig. 3d). In contrast, proteins enriched in LNR EVs demonstrate a distinct cellular stress and proteostasis signature at baseline (PSME1, PSME3, API5) (Fig. 3d). Collectively, the data reveal distinct protein signatures associated to BD neurons and Li responsiveness, which require further investigation for roles in BD diagnosis and/or pathophysiology.

### Differentially expressed miRNAs in EVs from patient-derived neuronal cultures

In addition to protein, we extracted and sequenced miRNA isolated from EVs. Of 417 identified miRNAs we found 7 miRNAs that were upregulated in BD neuronal EVs, and 6 miRNAs that were downregulated (Fig. 4a&c) relative to CTL neurons. Molecular function gene ontology (GO MF) demonstrated that the miRNA target genes related to DNA and RNA transcription factor binding, as well as protein kinase and phosphatase binding (Supp. Fig. S3a-c). Synaptic gene ontology (SynGO) results indicated that the differentially expressed microRNAs modulate genes related to synaptic transmission and post-synaptic function (Supp. Fig. S3a-c). We also identified 20 miRNAs differentially expressed between LR and LNR (Fig. 4b-c). GO MF analysis linked these miRNAs to DNA and RNA regulation (Supp. Fig. S4a-c). Additionally, downregulated miRNAs associated to cadherin, ephrin, and kinase binding were identified (Supp. Fig. S4a-c). SynGO results for miRNAs in LR were linked to multiple genes regulating neuronal post-synaptic membranes (Supp. Fig. S4a-c). Notably, 4 (CTL vs BD) and 6 (LR vs LNR) miRNAs that we identified as differentially expressed have been previously described in the literature^18, 34, 35^, reinforcing them as replicable candidates for biomarkers (Fig. 4d-e).

**Figure 4:**
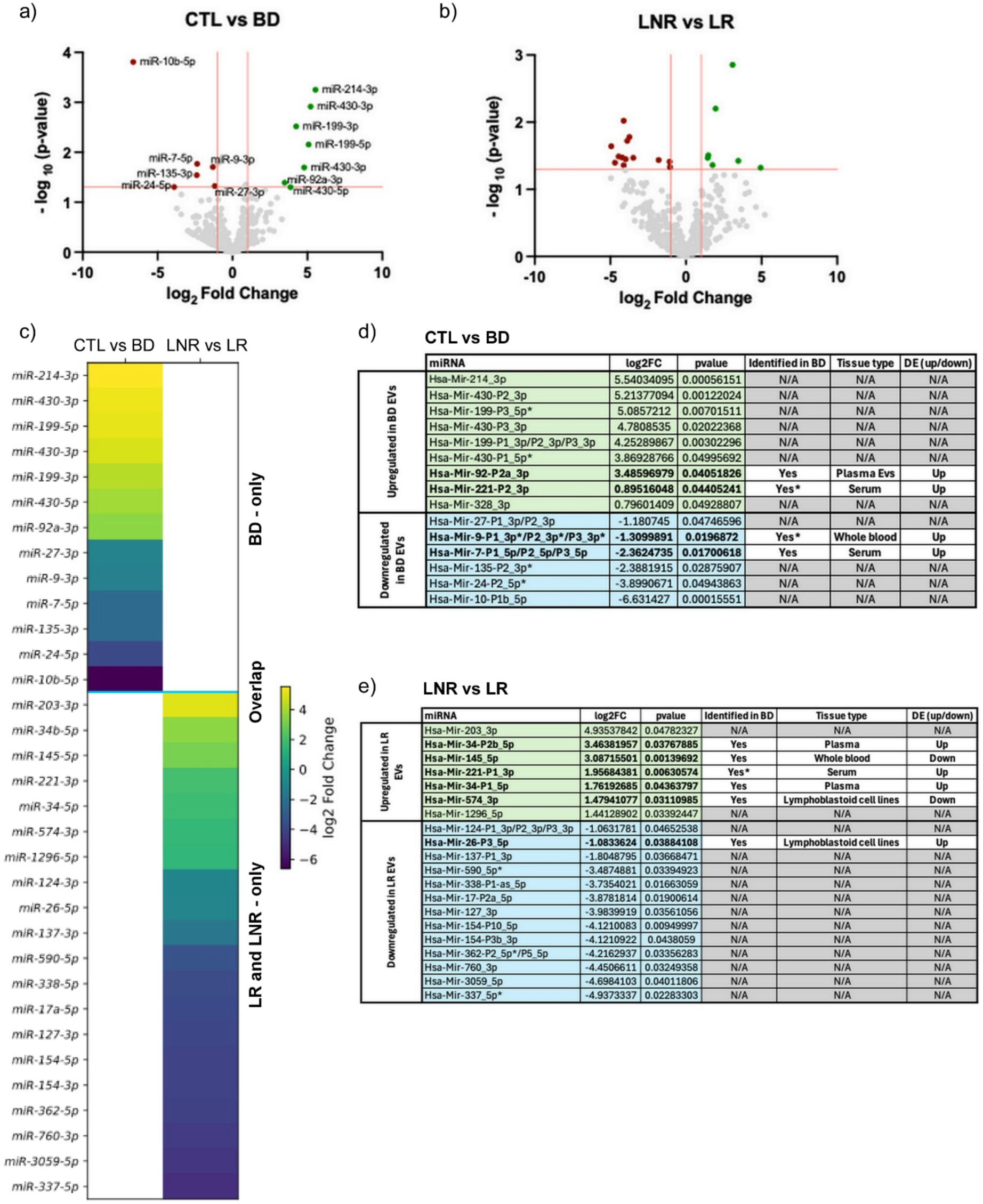
miRNA profiling of patient-derived EVs. **a)** Volcano plot comparing differential miRNA expression in EVs from CTL (n=3) vs BD (n=8) neuronal cultures. *X-intercept: Log_2_FC= ±1. Y-intercept: –log_10_(P-value) = 1.3 (p = 0.05). Points in grey are either not significant or falls outside fold change cutoff. Points in green are enriched In BD EV’s. Points in red are enriched in CTL EVs. **b)** Same as in **(a)** but for LNR (n=4) vs LR (n=4). **c)*** Heatmap of differentially expressed miRNAs from **(a)** & **(c). d-e)** Significantly upregulated (green) and downregulated (blue) miRNAs, their Log2FC, p-value, previous identification in BD literature [17,33,34], literature tissue origin, direction of expression change in the literature. **\*Yes:** indicates miRNAs of the same family that were identified, but not the specific arm. *Statistics: Wald’s tests with FDR<0.1%, p<0.05. Points in green are enriched In LR EV’s. Points in red are enriched in LNR EVs*.

Together, the results nominate candidate biomarkers for BD and Li responsiveness and provide insight into a mechanistic contribution of the neuronal secretome in BD pathogenesis.

### Assessing the effect of the BD neuronal secretome on network physiology

We hypothesized that the significant alterations to EV protein and miRNA cargo in BD neuronal secretions may influence inter-neuronal communication and play a role in the hyperexcitability phenotype we and others have observed^8–10^.

### Network hyperexcitability and altered response to potassium channel blockade

Utilizing the improved temporal resolution of GCaMP6f, expressed in neurons by AAV on a human synapsin promoter, we reproduced the hyperexcitability phenotype we previously observed by dye-based calcium imaging in BD neurons, relative to controls (Fig. 5a-c & Supp. Fig. S5a). Potassium leak channels set the rheostat for neuronal membrane excitability; their dysfunction has been implicated in BD^5, 8, 36^ which might underlie the hyperexcitability observed Thus to further interrogate this phenotype by maximizing release through elevated potassium channel function, we also assayed the effects of 4-aminopyridine (4-AP), a weak blocker of voltage-gated potassium channels. As expected, 4-AP significantly increased Ca^2+^ event frequency in CTL neurons (Fig. 5d & Supp. Fig. S5b), consistent with its known effect of prolonging depolarization and enhancing firing. However, BD neurons exhibited a modest reduction in firing following 4-AP (Fig. 5e & Supp. Fig. S5c-f) suggesting K+ channel flux in active BD neurons might be saturated. An increase in the proportion of active cells in both groups following blockade demonstrated that 4-AP treatment was effective (Fig. 5f-g & Supp. Fig. S5 g&h). The data support the conclusion that altered potassium channel function is present in BD neurons, but also that intrinsic (albeit insufficient) compensatory mechanisms are at play to limit further increases in network excitability.

**Figure 5:**
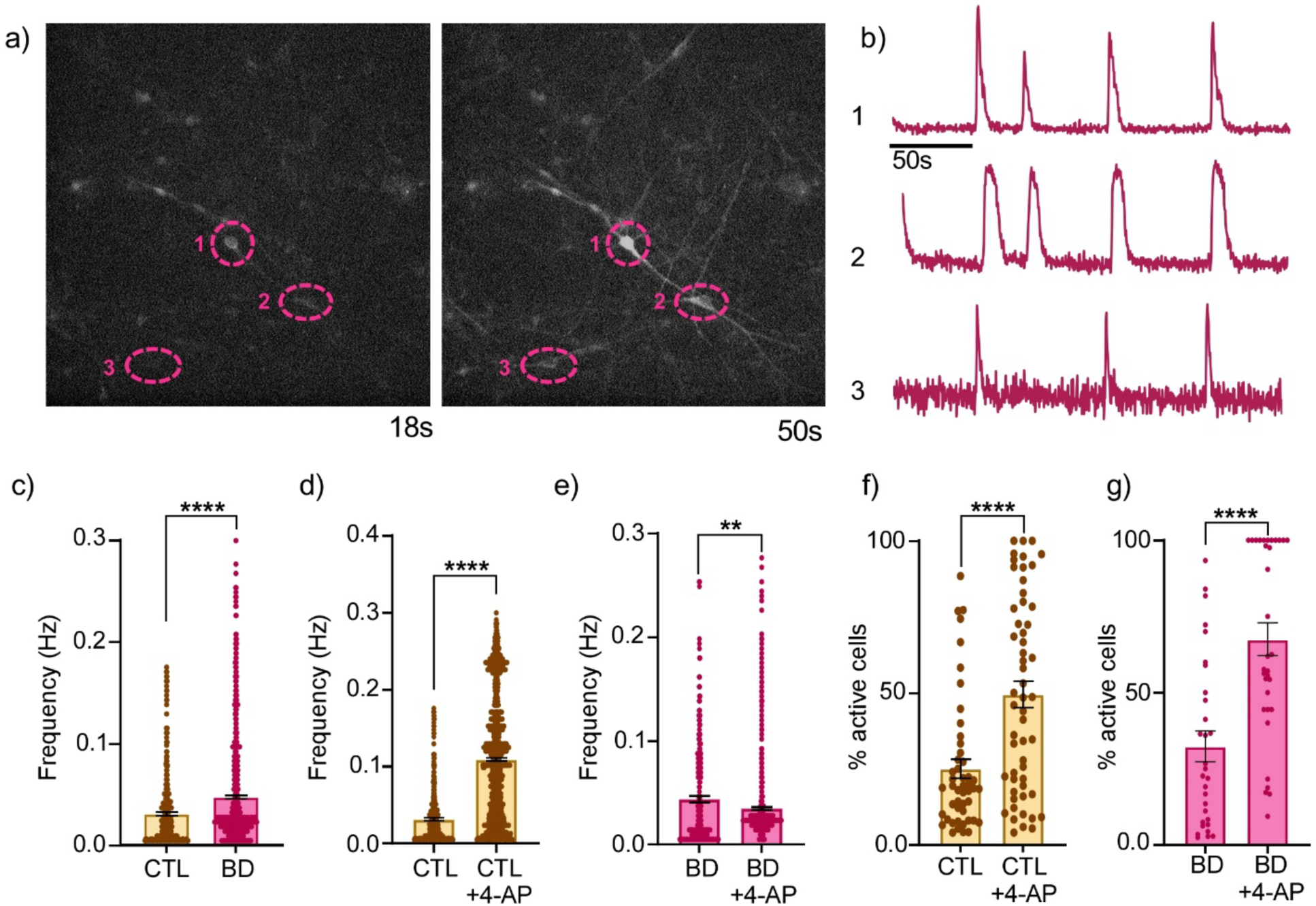
Replication of the BD hyperexcitability phenotype and characterization of 4-AP-induced modulation of neuronal network activity across control and BD lines. **a)** Representative timelapse of calcium imaging of iPSC-derived neurons using GCaMP6f. **b)** Representative raw fluorescence traces of spontaneous GCaMP6f calcium transients corresponding to the cell bodies (ROIs) denoted by pink circles in **(a)**. **c)** Calcium event frequency per neuron for CTL (n=3, 430 cells) and BD (LR & LNR, n=6, 1000 cells) neurons. **d)** Calcium event frequency per neuron before and after treatment with 4-aminopyridine (2.5 mM 4-AP) for CTL (n=3, 500-1200 cells) and **e)** BD (LR & LNR, n=6, 300-900 cells) neurons. **f)** Quantification of the percentage of active neurons per image field before and after treatment with 4-AP (2.5 mM) in CTL (n=3) and **g)** BD (LR & LNR, n=6) neurons. *Data shown are mean ±SEM. Statistics: two-tailed Welch’s t-test. **p < 0.005, ****p < 0.0001*.

### The BD secretome modulates neuronal network responses

To assess whether secreted material from BD neurons could contribute to network dysfunction, we conducted GCaMP6f recording in CTL neurons +/- chronic exposure to media conditioned with BD cultures (Fig. 6a). Under basal conditions, BD secretome exposure did not alter the proportion of active neurons (Fig. 6b) or calcium transient frequency (Fig 6c & Supp. Fig. S6a). Upon application of 4-AP, both groups showed the expected increase in network activity; however, CTL neurons exposed to BD-conditioned media exhibited a ∼30% greater increase in the proportion of active cells compared to untreated controls (Fig. 6b). In contrast, the 4-AP–induced increase in burst frequency was attenuated in BD-conditioned neurons (Fig. 6c; Supp. Fig. S6a), indicating an altered response to network stimulation. This suggests that the BD secretome engages mechanisms that attempt to regulate the underlying hyperexcitability observed in BD neuronal networks.

**Figure 6:**
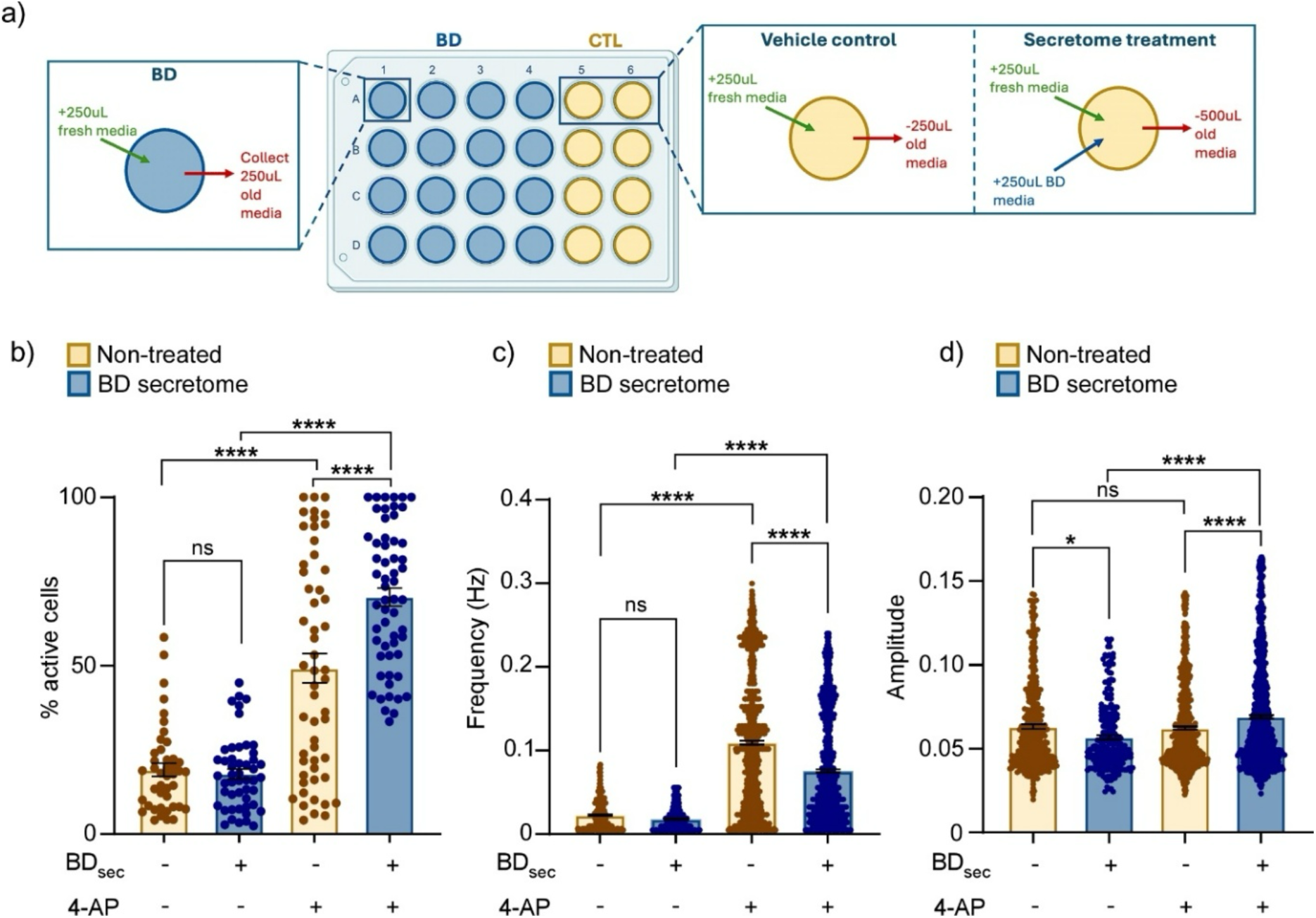
Using GCaMP6f to Investigate the effect of the BD secretome on CTL neuronal network activity. **a)** Schematic depicting neuronal secretome treatment pipeline. Media from BD patient-derived human neurons is collected and used to treat control patient-derived human neurons. *Vehicle control consists of a half-media change. BD media denotes conditioned media from BD coverslips (BD_sec_). Green and blue arrows indicate media being added; red arrows indicate media being removed/collected.* **b)** Quantification of the percentage of active neurons per image field in NT (yellow) and BD secretome (BD_sec_) treated (blue) CTL neurons before and after treatment with 4-AP (2.5 mM). **c)** Quantification of calcium event frequency per neuron and **d)** amplitude per neuron in NT (yellow) and BD secretome (BD_sec_) treated (blue) CTL neurons before and after treatment with 4-AP (2.5 mM). *Data shown are mean ±SEM. Statistics: Outliers were identified using the ROUT method (Q=1%). Statistical comparisons were performed using one-way ANOVA with Šídák’s multiple comparisons test. **\***p < 0.05, **\*\*\*\***p < 0.0001*.

At baseline, BD-conditioned neurons showed a modest but significant reduction in calcium transient amplitude, which was reversed under 4-AP, resulting in increased amplitude under conditions of maximal excitability (Fig. 6d; Supp. Fig. S6b). Similar patterns were observed for the area under the curve, integrating both amplitude and duration (Supp. Fig. S6c–d). Together, the data show that the BD secretome is bioactive, and enhances intrinsic excitability in control neurons, an effect that becomes apparent under conditions of maximal stimulation. The absence of this effect at baseline suggests that the network engages compensatory homeostatic mechanisms to dampen this increased excitatory drive.

## Discussion

This study identifies EV signatures associated with BD and lithium response using patient-derived iPSC neurons. We show BD neurons exhibit increased EV secretion, carry specific neuronal EV protein and miRNA signatures that regulate intrinsic network hyperexcitability. Importantly, the modulation of neuronal network responses suggesting a functional role for secreted factors in BD neural network dysfunction.

We observed increased EV release in LR neurons, with a similar but non-significant trend in LNR. While both groups are hyperexcitable, differences in EV secretion may reflect distinct underlying molecular mechanisms further strengthened by the leftward shift in EV size profile of LR neurons compared to LNR and CTL. Considering the literature on altered small GTPase signalling in LR neurons^8^, its modulation by lithium treatment^8^, and roles in intracellular trafficking pathways and EV biogenesis^37, 38^, further investigation of Li’s effect on the size and concentration of EVs released from BD neurons is warranted.

Proteomic analysis of EVs from BD patient neurons revealed 10 differentially expressed proteins relative to CTL. The most noteworthy being NRXN1 (neurexin-1) and KIDINS220 (Kinase D-interacting substrate of 220 kDa), as they are strong candidate biomarkers with a clear functional link to bipolar disorder phenomena. NRXN1 is a presynaptic cell adhesion molecule critical for synapse assembly and maturation that has been associated with several neuropsychiatric disorders including BD, SCZ, and autism spectrum disorder^39–41^. Since neurexins are enriched at presynaptic membranes, their presence in BD EVs may signify alterations in intercellular signalling involving synaptic communication or compensatory remodeling in BD.

KIDINS220 is a scaffolding protein and a downstream target of neurotrophic signalling in the CNS^42^ which plays a role in neuronal maturation, activity, and plasticity^42, 43^. This makes KIDINS220 a functionally relevant candidate biomarker because deficits in neurotrophic signalling, mainly concerning BDNF, have been heavily implicated in BD and other neuropsychiatric disorders^43^. Enrichment of KIDINS220 in BD EVs could point to disrupted neurotrophic signalling, in addition to modulation of synaptic activity and plasticity in BD neurons.

Interestingly, LIPG (endothelial lipase) and LPL (lipoprotein lipase), two proteins involved in lipid metabolism, were also enriched in BD EVs relative to CTL. This is particularly interesting considering the elevated levels of metabolic syndrome in BD patients^44^. It has been shown that individuals with comorbid BD and metabolic disorders such as type 2 diabetes (T2DM) and insulin resistance (IR) have a more severe prognosis and poorer treatment response^45^. In fact, metformin, an AMPK activator, and a first-line medication in the treatment of T2DM, has been shown to significantly improve symptoms of depression and anxiety in individuals with comorbid IR and treatment-resistant bipolar depression^46^. Enrichment of these metabolic proteins in BD neuronal EVs may reflect altered neuronal lipid metabolism, potentially linking BD neuropathology and metabolic risk.

We also compared EV content between LR and LNR to identify potential biomarkers to indicate beneficial treatment selection. The results suggest stark baseline biological differences between the two patient groups. While LR EVs were enriched with proteins involved in synaptic plasticity, neurotrophic signalling, and ion homeostasis, LNR EVs were enriched with proteins involved in stress responses and proteosome-related proteins. This contrast between patient neurons may indicate that lithium non-responsiveness may be associated with heightened cellular stress responses, rather than synaptic-related dysfunction seen in LR neurons.

The ATPase Na+/K+ transporting subunit alpha 3 (ATP1A3) was a particularly compelling hit in the LR EVs. The ATP1A3 isoform has been linked to BD^47, 48^ and described as a rescue pump for highly excitable neurons that undergo large increases in intracellular Na+ concentration^32^. Considering the hyperexcitability phenotype we observed in BD patient-neurons, this represents a biologically informative candidate biomarker for Li responsiveness.

Similarly, PTN (pleiotrophin), a secreted protein involved in neuron outgrowth, synaptic plasticity, and survival also emerged as a differentially expressed protein in LR. PTN has been shown to phosphorylate GSK3B, a direct target of Li treatment^49^ and was recently implicated in the regulation of insulin resistance^50^. Together, these differentially expressed proteins are strong biomarker candidates for lithium response, that require future validation in clinical patient biofluids.

In addition to candidate protein biomarkers, we also identified several EV miRNAs that may serve as molecular signatures of BD. Several EV-associated miRNAs identified in this study overlap with previously reported BD-associated miRNAs, supporting their potential utility as biomarkers. However, inconsistencies in directionality compared to peripheral studies highlight the importance of cellular context and EV-specific miRNA sorting. Although this study would have benefitted from a larger sample size (due to the variability of patient-derived hiPSC models) we have identified several candidate protein and miRNA biomarkers. We note that the differentially expressed miRNAs did not maintain significance after correction for multiple comparisons, but we provide several hits for validation of predictive use in patient biofluids. Together, the data suggest that EV miRNAs reflect disease-relevant regulatory pathways, and that validation in patient biofluids is required.

Consistent with prior studies, BD neurons exhibited hyperexcitability, alongside an altered response to potassium channel blockade. The reduced responsiveness to 4-AP, despite the increase in proportion of active neurons in the network, suggests disrupted potassium channel function and compensatory network mechanisms. These findings align with genetic^5^ and electrophysiological^51^ evidence implicating potassium channel dysregulation in BD and support a role for ion channel dysfunction in disease pathophysiology.

Since we identified significant changes in the BD EV proteome and miRNA contents related to synaptic modulation, we also investigated the functional implications of the BD secretome on neuronal activity. Importantly, we demonstrate that the BD secretome alters neuronal network responses, enhancing recruitment of active neurons while constraining firing frequency under stimulated conditions. This suggests that secreted factors, including EVs, may contribute to network-level homeostatic regulation. These findings introduce a previously underexplored mechanism by which BD-associated cellular phenotypes may propagate across neuronal networks.

The observed alterations in network activity were induced by conditioned media which includes a host of soluble non-vesicular secreted factors in addition to EVs. Thus, there may be novel drivers of network change and adaptation that were not revealed in EV samples. Future experiments comparing EV enrichment protocols with filtered EV-depleted conditioned media may prove enlightening. A more detailed characterization of these alterations would provide deeper insight into BD pathophysiology and may reveal novel targets for modulating neuronal hyperexcitability.

## Conclusion & summary

We have identified biologically relevant candidate protein and miRNA biomarkers for BD diagnosis. We have also identified differentially expressed proteins and miRNAs within EVs of LR and LNR, which may allow us to predict patient responses to Li. Several of the differentially expressed miRNAs we found were previously identified in patient biofluids in other studies, arguing for further validation in patient samples. Considering the excessive delay for diagnosis and treatment of BD patients, and increased suicide risk associated with this delay, our study provides a step toward expediting effective management and improving the quality of life for individuals with BD.

In agreement with the literature, our functional experiments have implicated potassium channel dysregulation in the now multiple-replicated BD hyperexcitability phenotype. Notably, we also demonstrated that the BD secretome influences neuronal network excitability and homeostasis in otherwise normal networks. To our knowledge, this is a first observation and provides a novel insight into the pathophysiology of BD. Gaining a deeper understanding of the functional implications of the BD secretome could uncover new therapeutic targets to regulate the hyperexcitability observed in BD patient-derived neurons.

## Authors Contributions

AP performed all cell culture with the help of YL, MAb, IP, and ZY. AP performed all experiments and calcium imaging. NTA was performed with the help of the Center for Applied Nanomedicine at the McGill University Health Center, with analysis done by AP. AP isolated the EVs, and analyzed the LC-MS/MS experiments. RNA extraction and sequencing was done by AP with the help of DR and Genome Quebec. Analysis of RNA extraction was done by CC in collaboration with AK and AP. Analysis script for GCaMP6f was written by CTR and conceptualized in collaboration with AP.

AP, AM and AK designed the study and made interpretations of the results; AP drafted the first version of the manuscript with inputs from AM and AK. GAR, MA, AM and AK provided supervision and funding.

All authors reviewed and edited the manuscript and approved the final version of the manuscript.

## Acknowledgements

We gratefully acknowledge the financial support of Bell Let’s Talk-Brain Canada Mental Research Program (AM, MA, & G.A.R) #5450, the Brain & Behavior Research Foundation Young Investigator Award (AK) #30822, and the Fonds de recherche du Québec - Santé (AP) #343031. We thank the Centre for Applied Nanomedicine (RI-MUHC), the center for Advanced Proteomic Analyses (IRIC), the Facility for Electron Microscopy at McGill, Génome Québec, and the Canadian Neurophotonics Platform Viral Vector Core Facility.

