## Supplemental Tables and Figures for "Neuronal secretome from bipolar patient-derived neurons alters network function and contains candidate biomarkers for diagnosis and lithium response"

| | Line number | Origin | Patient Ethnicity | Sex | Age | BD Diagnosis | Li Response |
|---|---|---|---|---|---|---|---|
| Controls | SBP009 | Lymphocyte | Caucasian | Male | 51 | n/a | n/a |
| | SBP012 | Lymphocyte | Caucasian | Male | 53 | n/a | n/a |
| | SBP008 | Lymphocyte | Caucasian | Male | 25 | n/a | n/a |
| | 3448 | PBMC | Caucasian | Male | 50 | n/a | n/a |
| | Kif4439 | PBMC | Caucasian | Male | 29 | n/a | n/a |
| Lithium Non-responders | SBP001 | Lymphocyte | Caucasian | Male | 51 | BD1 | 3/10 |
| | SBP002 | Lymphocyte | Caucasian | Male | 58 | BD1 | 1/10 |
| | SBP004 | Lymphocyte | Caucasian | Male | 40 | BD1 | 0/10 |
| | 4120 | PBMC | Caucasian | Male | 55 | BD1 | 0/10 |
| | 3980 | PBMC | Caucasian | Male | 30 | BD1 | 3/10 |
| Lithium Responders | SBP005 | Lymphocyte | Caucasian | Male | 41 | BD1 | 10/10 |
| | SBP007 | Lymphocyte | Caucasian | Male | 34 | BD1 | 9/10 |
| | SBP010 | Lymphocyte | Caucasian | Male | 50 | BD1 | 9/10 |
| | 4119 | PBMC | Caucasian | Male | 54 | BD1 | 9/10 |
| | 4146 | PBMC | Caucasian | Male | 37 | BD1 | 9/10 |

**Table S1: Clinical details of patient-derived IPSC lines.**

PBMC, peripheral blood mononuclear cells; BD1, Bipolar disorder type 1; Li response was determined via clinical assessment with the standardized Alda scale [1-3].

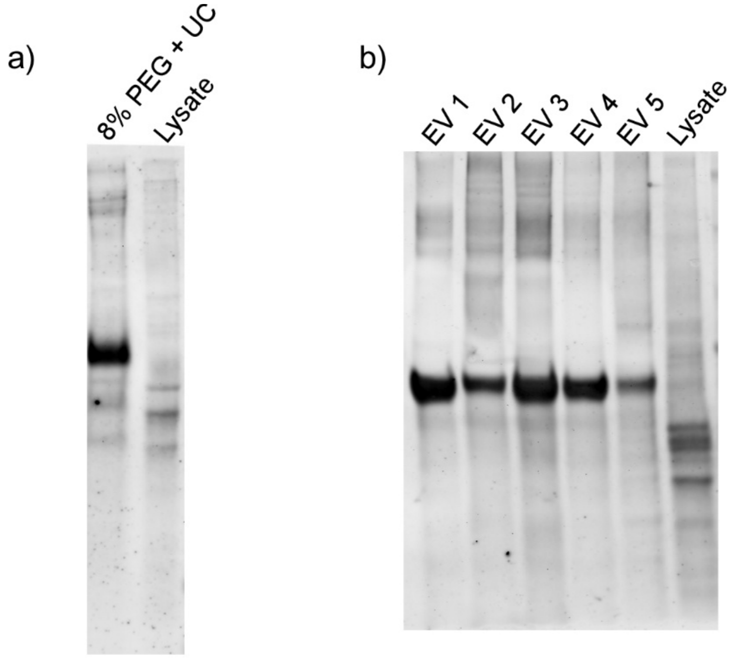


**Figure S1: Total protein stain for EV isolation characterization. a)** Total protein stain for western-blot from figure 1a. **b)** Total protein stain for western-blot from **Figure 1b.**

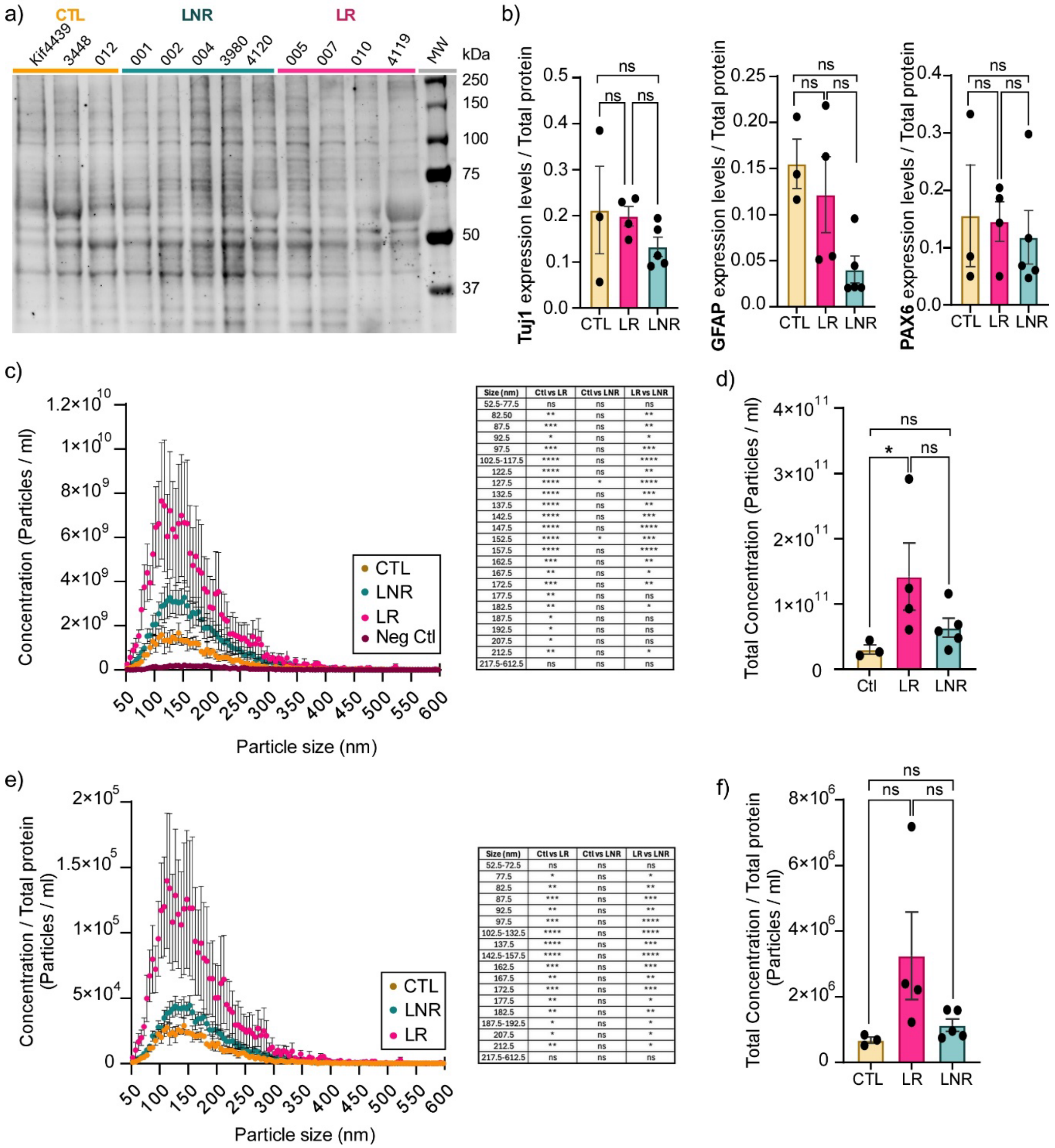


| Size (nm) | Ctl vs LR | Ctl vs LNR | LR vs LNR |
|---|---|---|---|
| 52.5-77.5 | ns | ns | ns |
| 82.50 | ** | ns | ** |
| 87.5 | *** | ns | ** |
| 92.5 | * | ns | * |
| 97.5 | *** | ns | *** |
| 102.5-117.5 | **** | ns | **** |
| 122.5 | **** | ns | ** |
| 127.5 | **** | * | **** |
| 132.5 | **** | ns | *** |
| 137.5 | **** | ns | ** |
| 142.5 | **** | ns | *** |
| 147.5 | **** | ns | **** |
| 152.5 | **** | * | *** |
| 157.5 | **** | ns | **** |
| 162.5 | *** | ns | ** |
| 167.5 | ** | ns | * |
| 172.5 | *** | ns | ** |
| 177.5 | ** | ns | ns |
| 182.5 | ** | ns | * |
| 187.5 | * | ns | ns |
| 192.5 | * | ns | ns |
| 207.5 | * | ns | ns |
| 212.5 | ** | ns | * |
| 217.5-612.5 | ns | ns | ns |

| Size (nm) | Ctl vs LR | Ctl vs LNR | LR vs LNR |
|---|---|---|---|
| 52.5-72.5 | ns | ns | ns |
| 77.5 | * | ns | * |
| 82.5 | ** | ns | ** |
| 87.5 | *** | ns | *** |
| 92.5 | ** | ns | ** |
| 97.5 | *** | ns | **** |
| 102.5-132.5 | **** | ns | **** |
| 137.5 | **** | ns | *** |
| 142.5-157.5 | **** | ns | **** |
| 162.5 | *** | ns | *** |
| 167.5 | ** | ns | ** |
| 172.5 | *** | ns | *** |
| 177.5 | ** | ns | * |
| 182.5 | ** | ns | ** |
| 187.5-192.5 | * | ns | * |
| 207.5 | * | ns | * |
| 212.5 | ** | ns | * |
| 217.5-612.5 | ns | ns | ns |

**Figure S2: Raw and total protein-normalized NTA results. a)** Total protein stain of all patient lines used for NTA. **b)** Normalization of neuronal (Tuj1), astrocytic (GFAP), and NPC (PAX6) composition from **Fig. 2a** to total protein stain in **a). c)** Raw NTA of isolated EVs from blank media (red, n=1), CTL (yellow, n=3), LR (pink, n=4), and LNR (turquoise, n=5). **d)** Quantification of total concentration of EVs from **c)**. **e)** NTA of isolated EVs from CTL (yellow, n=3), LR (pink, n=4), and LNR (turquoise, n=5) normalized to total protein. **f)** Quantification of total concentration of EVs from **e).** *Data shown are mean ±SEM. Statistics: One-way ANOVA* ***(d&f)*** *and two-way ANOVA* ***(c&e)*** *with Holm-Šídák's multiple comparisons test was used. *p<0.0332, **p<0.0021, ***p<0.0002, ****p<0.0001.*

a) CTL vs BD

| hsa-miR-214-3p | hsa-miR-221-3p | | hsa-miR-92a-3p | hsa-miR-328-3p |
|---|---|---|---|---|
| EZH2 | CDKN1B | CTCF | ITGA5 | ABCG2 |
| PTEN | BMF | RAB1A | TP63 | CD44 |
| MAP2K3 | FOXO3 | RECK | BMPR2 | H2AFX |
| MAPK8 | KIT | SIRT1 | BCL2L11 | MMP16 |
| POU4F2 | CDKN1C | MDM2 | STAT3 | CPT1A |
| PLXNB1 | TMED7 | SOCS1 | HIPK1 | NRBP1 |
| SRGAP1 | DDIT4 | ARF4 | NR1H4 | |
| XBP1 | BNIP3L | CXCL12 | PTEN | |
| TWIST1 | BBC3 | ADAMTS6 | SOCS5 | |
| CTNNB1 | TIMP3 | CREB1 | MAPK8 | |
| BCL2L2 | FOS | GAS5 | MAP2K4 | |
| ARL 2.00 | BNIP3 | TP53INP1 | DUSP10 | |
| JAG1 | ESR1 | HMGA1 | CCL8 | |
| UBE2I | TICAM1 | SOCS5 | CD69 | |
| FGFR1 | PTEN | HIF1A | FBXW7 | |
| NRAS | DIRAS3 | BRWD3 | LATS2 | |
| BCL2L11 | ETS1 | | NRF1 | |
| CDK6 | TRPS1 | | MTO1 | |
| ALPK2 | CERS2 | | WWOX | |
| PAPPA | WEE1 | | TF | |
| LZTS1 | HECTD2 | | DCST1-AS1 | |
| MEF2C | ZEB2 | | | |
| IKBKB | ASZ1 | | | |
| ABCB1 | PDGFA | | | |
| VEGFA | RB1 | | | |
| BCL2 | APAF1 | | | |
| SNHG3 | ANXA1 | | | |

| hsa-miR-7-5p | hsa-miR-30a-5p | hsa-miR-10b-5p | hsa-miR-9-3p |
|---|---|---|---|
| SNCA | NOTCH1 | HOXD10 | EIF5A2 |
| EGFR | BECN1 | KLF4 | EGFR |
| RAF1 | DTL | PPARA | |
| CKAP4 | SNAI1 | TIAM1 | |
| PSME3 | PIK3CD | HTATIP2 | |
| SRSF1 | FOXD1 | | |
| SLC7A5 | KIF11 | | |
| IGF1R | VIM | | |
| PTK2 | RUNX2 | | |
| HOXB5 | ERG | | |
| BCL2 | BCL9 | | |
| OSBPL11 | EYA2 | | |
| PIK3R3 | SOX4 | | |
| XRCC2 | CBX3 | | |
| KLF4 | TP53 | | |
| CCNE1 | CD99 | | |
| CUL5 | MAPK8 | | |
| SERPINB5 | NCAM1 | | |
| FOS | ATF1 | | |
| REL | ATG5 | | |
| GDF5 | | | |
| AR | | | |
| NOTCH4 | | | |
| Sp1 | | | |
| LINC00240 | | | |
| RAPGEF3 | | | |

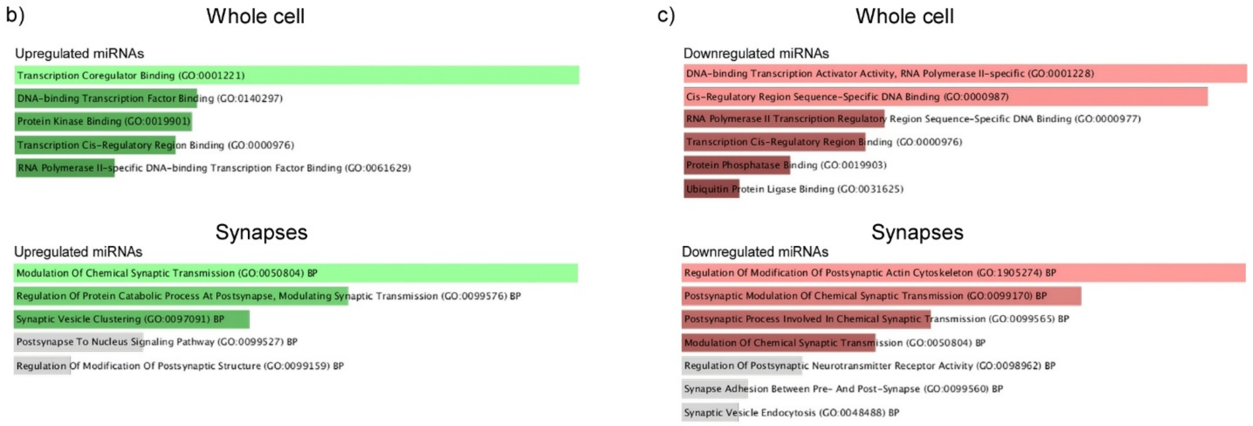


**Figure S3: Experimentally validated gene targets for differentially expressed miRNAs in CTL vs BD EVs determined using miRTarBase [40]. a)** experimentally validated gene targets for upregulated (green) and downregulated (red) miRNAs in BD (n=8) EVs relative to CTL (n=3). **b)** GO molecular function and Synaptic GO terms for gene targets of upregulated (green) **(b)** and downregulated(red) **(c)** miRNAs shown in **a)**. *GO terms are ranked by adjusted p-value. Terms in grey are not significant.*

a) LNR vs LR

| hsa-miR-145-5p | | hsa-miR-221-3p | | hsa-miR-574-3p | hsa-miR-34b-5p | hsa-miR-1296-5p |
|---|---|---|---|---|---|---|
| BNIP3 | EPAS1 | CDKN1B | SOCS1 | RAC1 | BCL2 | SP1 |
| SOX2 | ETS1 | BMF | ARF4 | EGFR | CREB1 | |
| KLF4 | RREB1 | FOXO3 | CXCL12 | EP300 | TGFBR2 | |
| MUC1 | CD44 | KIT | ADAMTS6 | RXRA | | |
| MYO6 | BRAF | CDKN1C | CREB1 | SMAD4 | | |
| CDKN1A | SOX9 | TMED7 | GAS5 | Ccnd2 | | |
| STAT1 | SMAD3 | DDIT4 | TP53INP1 | ND5 | | |
| YES1 | TGFBR2 | BNIP3L | HMGA1 | | | |
| IRS1 | SMAD4 | BBC3 | SOCS5 | | | |
| HOXA9 | CTNND1 | TIMP3 | HIF1A | | | |
| FSCN1 | SP1 | FOS | BRWD3 | | | |
| MYC | TNFSF13 | BNIP3 | | | | |
| FLI1 | CDK6 | ESR1 | | | | |
| DFFA | DDX6 | TICAM1 | | | | |
| IFNB1 | ARF6 | PTEN | | | | |
| POU5F1 | ADD3 | DIRAS3 | | | | |
| IGF1R | HMGA2 | ETS1 | | | | |
| ROBO2 | E2F3 | TRPS1 | | | | |
| EIF4E | SP7 | CERS2 | | | | |
| CDK4 | HDAC11 | WEE1 | | | | |
| SERPINE1 | SENP1 | HECTD2 | | | | |
| ITGB8 | NAIP | ZEB2 | | | | |
| SWAP70 | TBX15 | ASZ1 | | | | |
| JAG1 | WNT2B | PDGFA | | | | |
| NEDD9 | FGF10 | RB1 | | | | |
| PAK4 | DNMT3a | APAF1 | | | | |
| DDX17 | SCHLAP1 | ANXA1 | | | | |
| NRAS | EZH2 | CTCF | | | | |
| ILK | NKX2-1-AS1 | RAB1A | | | | |
| ADAM17 | NOTCH1 | RECK | | | | |
| RTKN | IGF1 | SIRT1 | | | | |
| ABRACL | | MDM2 | | | | |

| hsa-miR-124-3p | | hsa-miR-590-5p | hsa-miR-760 | hsa-miR-154-5p | hsa-miR-338-5p |
|---|---|---|---|---|---|
| EFNB1 | SPHK1 | TGFBR2 | CSNK2A1 | TLR2 | LRP1 |
| MYH9 | TPST2 | CHL1 | HIST1H3D | HMGA2 | ID1 |
| CDK6 | AKT3 | PDCD4 | HIST1H2AD | CUL2 | |
| PTBP1 | RAB32 | CREB5 | ATXN1 | | |
| SLC16A1 | TFAP4 | STAT3 | ETS1 | | |
| B4GALT1 | FSTL1 | FTX | | | |
| IQGAP1 | CD151 | RBPJ | | | |
| CAV1 | RCAN1 | | | | |
| ITGB1 | PLEC | | | | |
| NFATC1 | TRIB3 | | | | |
| RHOG | STAT3 | | | | |
| PPP1R13L | JAG1 | | | | |
| CCL2 | CAMTA1 | | | | |
| VIM | CLOCK | | | | |
| SMYD3 | SYCP1 | | | | |
| E2F6 | HNRNPA2B1 | | | | |
| NFKBIZ | Rab38 | | | | |
| AR | AMOTL1 | | | | |
| ROCK2 | SOS 1 | | | | |
| IL6R | HOTAIR | | | | |
| HMGA1 | CD274 | | | | |
| ROCK1 | CBL | | | | |
| RRAS | LINC00240 | | | | |
| PRRX1 | ACTN4 | | | | |
| GRB2 | ABCC4 | | | | |
| RAB27A | CRKL | | | | |

b)

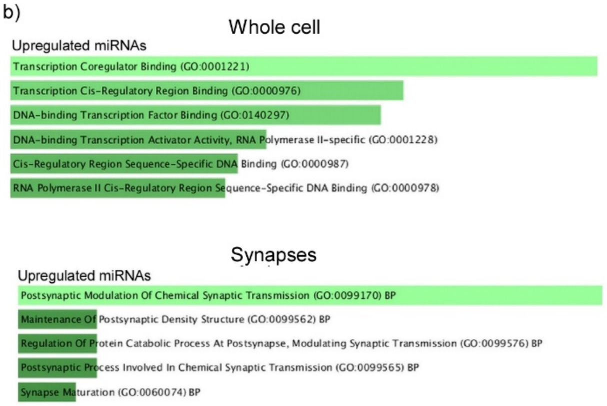


c)

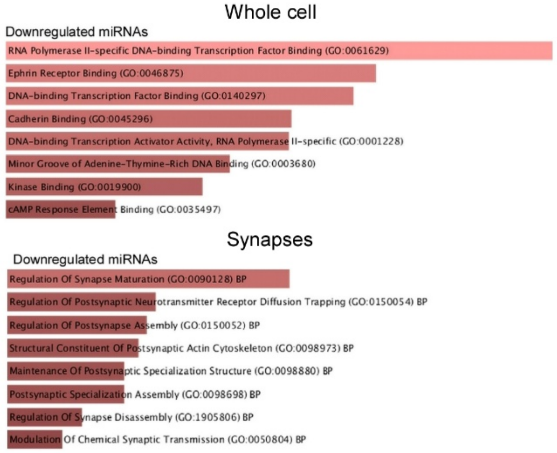


**Figure S4: Experimentally validated gene targets for differentially expressed miRNAs in LR vs LNR EVs determined using miRTarBase [40]. a)** experimentally validated gene targets for upregulated (green) and downregulated (red) miRNAs in LR (n=4) EVs relative to LNR (n=4). **b)** GO molecular function and Synaptic GO terms for gene targets of upregulated (green) **(b)** and downregulated(red) **(c)** miRNAs shown in **a)**. *GO terms are ranked by adjusted p-value.*

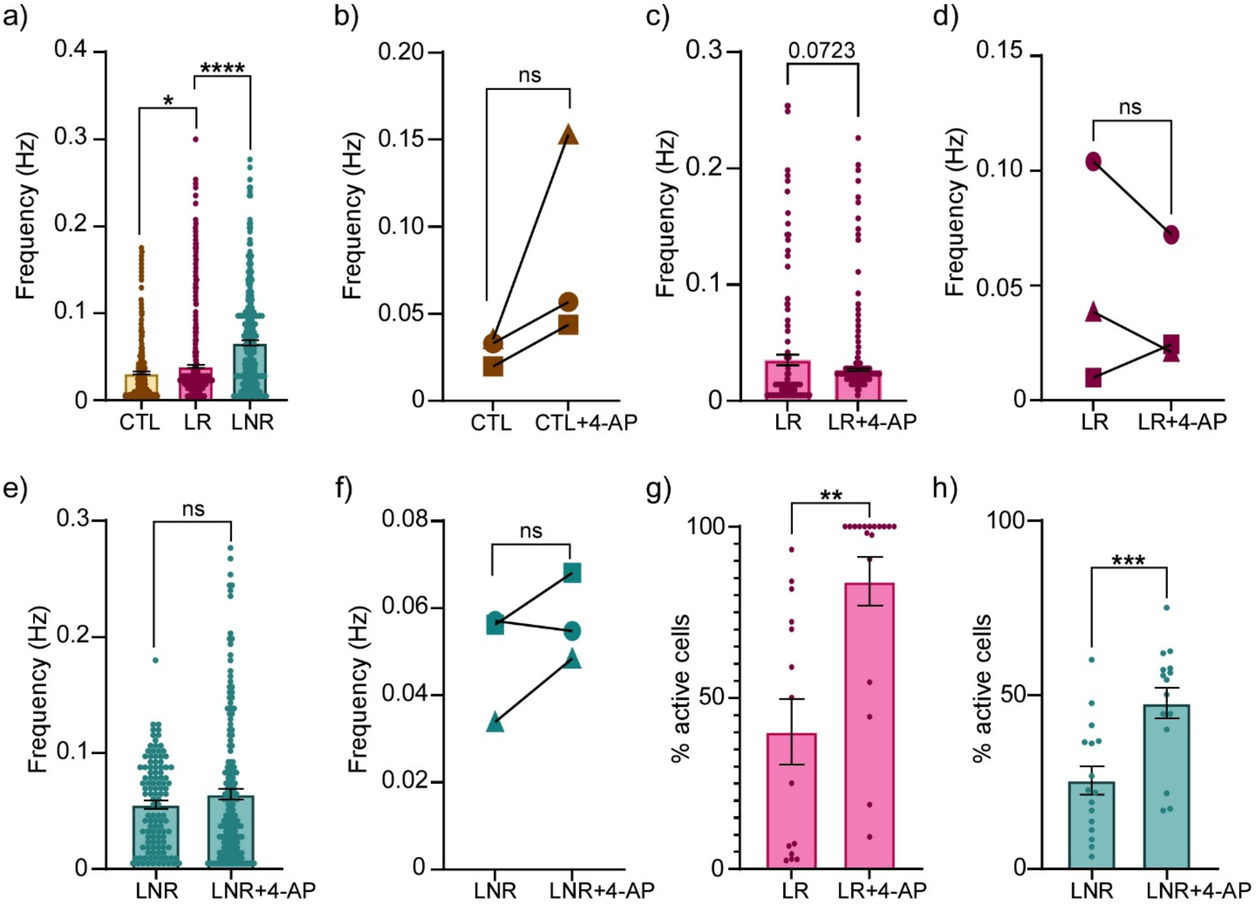


**Figure S5: Replication of the BD hyperexcitability phenotype and characterization of 4-AP-induced modulation of neuronal network activity across CTL, LR, and LNR lines. a)** Calcium event frequency per neuron for CTL (n=3, 426 cells), LR (n=3, 727 cells), and LNR (n=3, 343 cells). **b)** Calcium event frequency of CTL (n=3) per line before (n=426 cells) and after (n=1236 cells) treatment with 4-aminopyridine (2.5 mM 4-AP). **c)** Calcium event frequency of LR (n=3) per neuron and **d)** per line before (n=155 cells) and after (n=665 cells) treatment with 4-aminopyridine (2.5 mM 4-AP). **e)** Calcium event frequency of LNR (n=3) per neuron and **f)** per line before (n=113 cells) and after (n=179 cells) treatment with 4-aminopyridine (2.5 mM 4-AP). **g)** Percentage of active neurons per image field before and after treatment with 4-AP (2.5 mM) in LR (n=3), and **h)** LNR (n=3). *Data shown are mean ±SEM. Statistics:* ***a)*** *one-way ANOVA with Šídák's multiple comparisons test. *p < 0.05, ****p < 0.0001.* ***b-i)*** *Welch's t-test. **p<0.0021, ***p<0002*

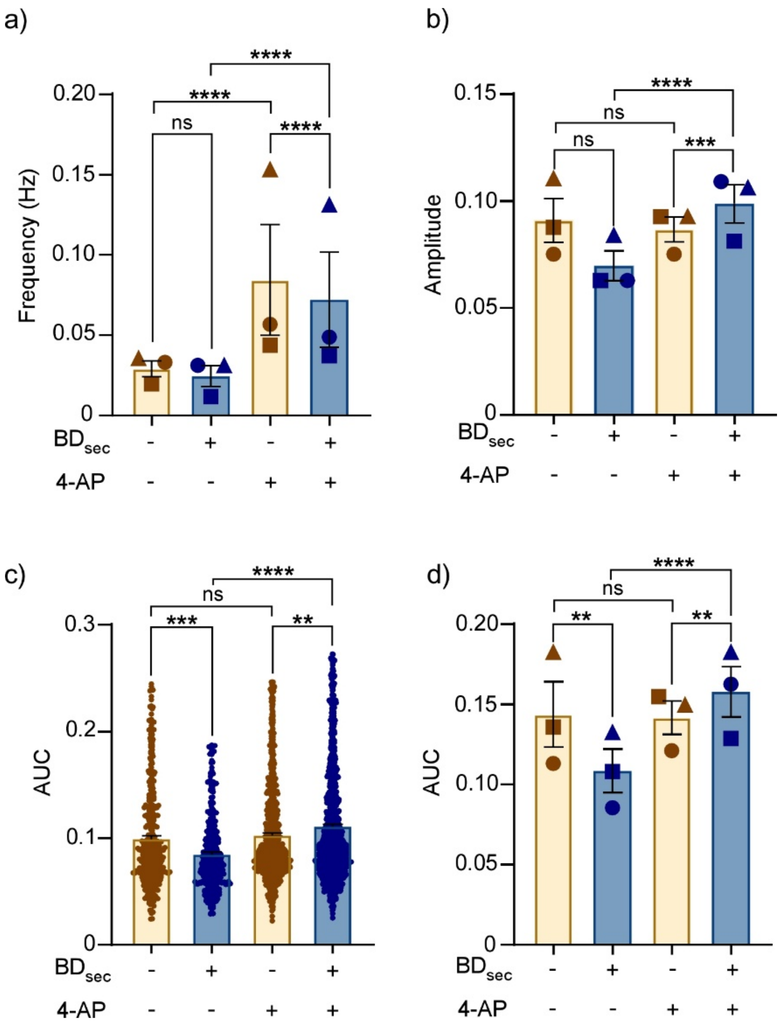


**Figure S6: Using GCaMP6f to Investigate the effect of the BD secretome on CTL neuronal network activity. a)** Calcium event frequency and **b)** amplitude per line (n=3, 2862 cells) in NT (yellow) and BD conditioned media ($BD_{cm}$) treated (blue) CTL neurons before and after treatment with 4-AP (2.5 mM). **c)** Quantification of area under the curve (AUC) per neuron and **d)** per line (n=3, 2862 cells) in NT (yellow) and BD conditioned media ($BD_{cm}$) treated (blue) CTL neurons before and after treatment with 4-AP (2.5 mM). *Data shown are mean ±SEM. Statistics: Statistical comparisons were performed using two-way ANOVA with Holm-Šídák's multiple comparisons test. **p < 0.0021, ***p < 0.0002, ****p<0.0001.*
